# Heavily burned wood from wildfires is less likely to provide functionality in streams

**DOI:** 10.1101/2020.10.25.354217

**Authors:** Pedro Gonçalves Vaz, Eric C. Merten, Christopher T. Robinson, Paulo Pinto

## Abstract

1. Increasingly severe forest fires are recruiting more heavily burned wood into streams. Wood affects every ecological and physical process in streams differently throughout seasons. However, little is known about the seasonality of wood functions in fire-prone biomes and how it combines with wood burning level to guide future postfire restoration efforts.
2. Through an extensive three-year seasonal tracking of stream wood following forest fires in central Portugal, we examined for the first time the influence of burning level, season, and a large suite of driving factors on the likelihood of each of four functions with primary ecological consequences — retention of organic matter, serving as substrate for aquatic biota, being key pieces forming wood jams, and deflecting flow including pool habitat formation.
3. Our results strongly support that one of the main ecological functions of wood in rivers, i.e. to provide substrate for biological organisms — namely for vegetation, periphyton, biofilms, and ovipositions — can be negatively affected in heavily burned wood.
4. Except for jam formation, the probability of each stream wood function changed markedly with season and the probability of non-function was nearly twice as high in the Euro-Mediterranean dry as in the wet season.
5. More anchored and decayed wood increased the probability of all functions, whereas the effect of submergence depended on the function. Challenging the “size paradigm” assuming larger-sized pieces to provide more function, our data suggest the effect of size to be function-specific.
6. *Synthesis and applications*. We show how postfire restoration success can be maximized by selecting the most appropriate wood, taking advantage of attribute-function relationships and choosing the right timing for operations. We urge managers to refrain from removing wood or to selectively remove the most heavily carbonized only, allowing the persistence of great potential to provide substrate for stream biota. The non-attraction of heavily burned wood as substrate can be compensated for by other wood with attributes enhancing this function, such as wood deeper within the bankfull area, and with large diameters. These results help to inform successful management, as is increasingly asked from restoration ecology.

## Introduction

Wood can affect virtually every biological and physical process in streams (Gregory et al. 2003, Coe et al. 2009, Merten et al. 2013, Molokwu et al. 2014). We refer to such wood pieces as functional; i.e., performing some observable function in the stream (Cordova et al. 2007, Vaz et al. 2013b). Functions with primary ecological consequences on streams include the retention of organic matter (Osei et al. 2015), serving as substrate for aquatic biota (McLachlan 1970), being key stable pieces forming wood jams (Abbe & Montgomery 2003) and deflecting flow (Mutz 2000). These wood functions have been examined globally for decades, but much less is known about what characteristics of individual wood pieces favor a particular function (Rosenfeld & Huato 2003). It is widely assumed that larger-sized and stable pieces provide more function. Yet, most studies have focused on the relationship between wood quantity and channel structure (Chen et al. 2008, Grabowski et al. 2019). In particular, it is not known how wood burning influences particular functions.

Burned wood may be straighter and have fewer branches than non-burned wood (Agee 1993), which may negatively affect its retention of matter, jam formation, and flow deflection functions. Burned wood may also be thicker in diameter and more decayed than non-burned wood, which may positively affect these aforementioned functions (Jones et al. 2011, Vaz et al. 2011). Fire-derived changes in physical and chemical properties may also affect wood substrate function. Water and extractants (namely lipids and terpenoid hydrocarbons) are lost in burned wood and charcoal tends to be biologically inert, whereas volatilization of repellent compounds may occur (Hyde et al. 2011), making burned wood arguably more attractive than unburned wood to provide substrate for organisms and ovipositioning (Vaz et al. 2014). Across streams with seasonal flow patterns in fire-prone biomes, seasonal drying (Verkaik et al. 2013) may play a role as important as burn status in wood functions (Flores et al. 2017). In wet seasons like spring in the Euro-Mediterranean region, wood functions such as matter retention, substrate provisioning, and flow deflection can be more enhanced than in the dry season with low or intermittent flows. The study of the effect of the burn status and season on functions will help practitioners seeking postfire restoration approaches that ‘work with natural processes’ to deliver ecological and geomorphological outcomes (Grabowski et al. 2019).

Incorporating wood into river restoration and management involves considering a set of wood attributes that may greatly alter function. Structural attributes include piece diameter, length, complexity (sensu Newbrey et al. 2005), decay state, and form (Cordova et al. 2007). For example, probability of flow deflection, including pool habitat formation, can increase with wood diameter (Magilligan et al. 2008). Longer and more complex wood can form jams (Abbe & Montgomery 2003). Decayed wood can contribute more to matter retention, jams, and flow deflection forming riffles and pools (Jones et al., 2011). Critical attributes concerning wood relationships with the stream channel include the level of submergence and how it rests within the channel (position), degree of anchoring, percentage within bankfull, and distance to the bank (Parsons & Thoms 2007). For instance, more submerged wood is more likely to serve as substrate for aquatic biota. More anchored and decayed wood, as indicators of stability and longevity, are relevant for a broad spectrum of wood functions. Although a great deal is known about how wood functions in rivers, these feature-function relationships have seldom been analyzed in a single dataset. Once assessed, these relationships can be harnessed to deliver multiple ecological benefits to the lotic ecosystem.

In this study, we analyzed the potential of combining wood burned level, season, and a large suite of functional driving factors to guide postfire restoration efforts in lotic ecosystems. To this end, we monitored over three years the influence of these factors on the likelihood of each of four primary functions — matter retention, substrate provisioning, jam formation, and flow deflection — by wood pieces in streams of burnt areas in central

Portugal. Because there seems to be a trend for large wood in rivers to become scarcer in these fire-prone forest areas (Silva et al. 2011, Moreira et al. 2011, Vaz et al. 2011, 2013a), each piece of wood will have increased importance and the maintenance of more functional wood in rivers is an important conservation target in these areas. Specifically, we addressed the following three questions: (a) do forest fires change the probabilities of specific stream wood functions through the burned level inflicted? (b) are functions stable between dry and wet seasons in a region like the Mediterranean or does the probability of each function change intra-annually? (c) how can well-documented wood structural attributes and those concerning its relationships with the channel affect specific wood functions together with the burned level?

## Materials and methods

### Study area and site selection

We conducted the study in central Portugal from early October 2010 to early May 2012 in a sub-basin of the Tagus River (Rio Frio) that experienced wildfires (71% burned area) between 2003 and 2007 (Fig.1). The local climate is Mediterranean with hot, dry summers and cool, wet winters. Mean annual precipitation is 512 mm (range: 3 mm in July to 82 mm in November) and mean annual temperature is 15.8 °C (range: 9 °C in December–January to 23 °C in July–August). Rio Frio (drainage area 37 km^2^, mean stream gradient 5.1 %) has gentle relief with altitudes ranging from 25 to 434 m (mean ~219 m). Geology at the streams was mainly characterized by siliceous rocks with low mineralization. Land cover was 55 % forest, 22 % shrublands, and 22 % agriculture. The dominant forest was maritime pine (*Pinus pinaster*), the species most affected by wildfires in Portugal (Moreira et al. 2009, Silva et al. 2009). In the study area, maritime pine is grown for timber in privately owned monoculture natural regeneration stands with a rotation of ≥ 35 years. In the most accessible areas (e.g., <25% slope), thinning to decrease tree density to ~235 stems/ha and shrub clearing were the most common management practices, generally excluding areas nearby streams.

**Figure 1.**
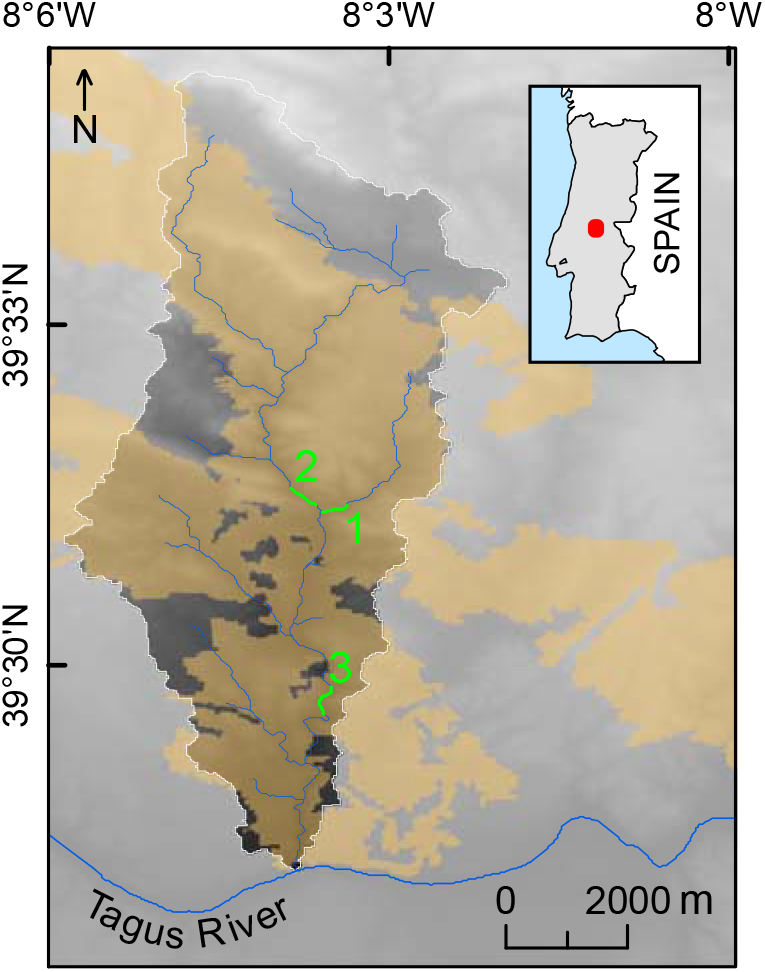
Location of the sampling sites (highlighted green lines) and 2003–2007 fire areas (orange polygons) in east-central Portugal. Three stream reaches were assessed (one each from 1^st^-, 2^nd^-, and 3^rd^-order streams; numbers stand for the orders) within the Rio Frio subbasin (white outlined) of the Tagus River.

Within Rio Frio, we selected three homogeneous reaches (~400 m each) having a burned sideband of at least 100 m (Vaz et al. 2013b), one each from stream order 1–3 (Strahler, 1957). Fire severity across the three stream reaches was similar, as visually and qualitatively assessed (e.g., tree mortality and amount of char in trunks). In the sideband, intense wildfires killed most pine trees (mostly <10 years old), and the dead trees fell within 1 to 3 years (PGV, personal observation). The understory was shrubby with dense growth of *Erica* spp., *Cistus* spp., and *Ulex* spp. and young postfire maritime pines. In the riparian zones, most trees did not die from fire. The dominant substrate was gravel with some boulders in the main channel. Mean channel widths were 2.9, 5.1, and 6.0 m in the first, second, and third order reaches, respectively. The reaches had neutral–basic waters and were intermittent, with stretches remaining dry for several months, in alternating dry and wet seasons. The wet season in the Euro-Mediterranean region can vary (e.g., Craveiro et al. 2019), but in the years of data collection it lasted from November to May. The natural discharge regime is primarily precipitation-dominated with highest discharge occurring in winter. Discharge responds rapidly to precipitation events (Fig. 2), which can result in major changes in flow over relatively short periods of time. Riparian zones with a distinct riparian community extended 5–15 m from the streams. The uncultivated riparian vegetation was dominated by ash (*Fraxinus angustifolia*), alder (*Alnus glatinosa*), black poplar (*Populus nigra*), and silver wattle (*Acacia dealbata*) with a few edges of bramble-thicket (*Rubus ulmifolius*). Some postfire logging was carried out outside the riparian zones but not nearby the streams. Fire-killed trees were left on the ground on stream side-slopes. Percent contributions from the adjacent pine forest for total instream wood volumes were ~25% and ~5% in the 1^st^–2^nd^ and 3rd order reaches, respectively (Vaz et al. 2013a).

**Figure 2.**
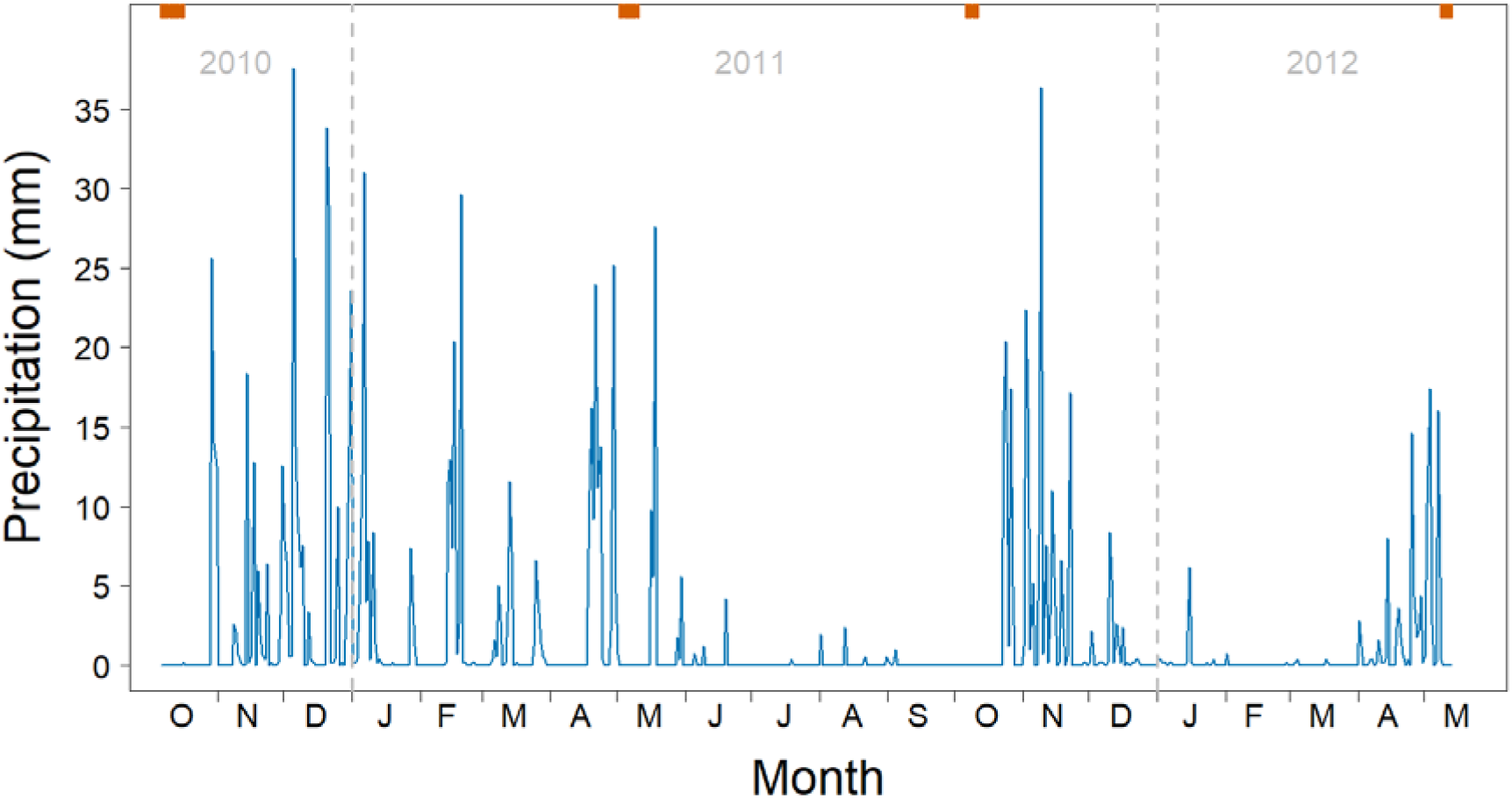
Local daily precipitation for the duration of the experiment. The rain gauge is 15 km from Rio Frio. Upper axis tick marks = data collection days.

### Data collection

To assess the intra-annual change in wood functions, we collected data on the three reaches in two dry seasons — early fall (6–16 October 2010, 6–11 October 2011) — and two wet seasons — late spring (2–10 May 2011, 8–13 May 2012). During the four surveys, we measured dead, downed wood pieces (diameter ≥ 0.05 m; length ≥ 0.5 m) and those that were still alive but entirely uprooted. We excluded snags, defined as pieces leaning or suspended over the stream at an angle greater than 30°. In wood jams (>2 pieces), we measured pieces that were accessible and whose functions were not influenced by the functions of other pieces. Only downed stream wood extending within bankfull boundaries were included in the tallies (Vaz et al. 2013b). To track wood characteristics and function, each piece was individually tagged, measured, and remeasured in the following season. We used one round blue pre-numbered anodized aluminum tag (32 mm diameter) on each end of each piece secured with a galvanized nail. For easier detection, we marked the tag place with white plastic-coated wire attached around the wood perimeter. The center of each piece was geo-referenced with a GPS unit (whenever possible, with a 0.3–1 m precision by post-processing).

Per each survey, we recorded the main function of the piece of wood in the channel regarding Retention, Substrate, Jams, or Flow (Table 1). We then recorded the following piece characteristics:

a. Wood burn level, assessed using three classes (unburned: no char; moderately burned: charred bark but outermost ring present in at least one part of the circumference; heavily burned: charred bark and sapwood resulting in significant ring loss, Jones & Daniels 2008, Vaz et al. 2013b);
b. Submergence, using three classes (vertically traversing the bankfull height, and lower or upper channel halves of the bankfull height);
c. Decay, using the four classes proposed by Jones and Daniels (2008) (evaluating bark, branches, and overall structural integrity), later simplified into two classes (sound, decayed), coalescing the first three to get balanced classes;
d. Form (straight, bent);
e. Position on the stream (ramp: resting on one bank only; bridge: log spans channel, touching both banks and resting on the floodplain; loose: resting entirely on the streambed). Due to a small sample number, bridges were coalesced with ramps for analysis;
f. Diameter, determined to the nearest 0.5 cm using a meter tape by a single measurement taken from a point considered the mean diameter by visual assessment;
g. Length, in meters to the nearest 0.01 m for the segments of the pieces that were >1 cm in diameter;
h. Percentage within bankfull; i.e., percentage part of the piece contained in the channel until the bankfull height;
i. Number of anchor ends; i.e., ends or sides attached (pinned under rocks, pinned under larger logs, or in channel spanning jams) or buried (in streambed sediment) in either the bank or the stream;
j. distance to bank; i.e., to the closest bank, in meters to the nearest 0.1 m; and complexity; i.e., branching complexity by counting attached branches and twigs according to Newbrey et al. (2005).

**Table 1.** Characterization of the wood functions monitored over three years in Portuguese streams.

| Name | Description |
| --- | --- |
| None | No function detected after careful observation of the whole wood piece remaining partly or fully within the bankfull channel |
| Retention | Retaining organic matter such as twigs, leaves, fine organic matter (observable volume $> \sim 10^{-3} \text{ m}^3$) |
| Substrate | Serving as a substrate for aquatic vegetation, periphyton, and/or epixylic biofilms (submersed wood with a conspicuous biofilm layer) or to conspicuous ovipositions (e.g., amphibians) |
| Jam | Creating debris jams (e.g., bracing other stream wood debris or serving as a key piece forming wood jams) |
| Flow | Deflecting flow (e.g., creating pools or riffles, forming steps). |

### Statistical Analysis

To evaluate the effects of wood burned level and season on probability of each stream wood function after fire, we used multinomial mixed-effects modeling with logit link conducted in a Bayesian framework. The multinomial response included the four wood functions (Retention, Substrate, Jam, and Flow) and None as the reference category. The fixed effects included the categorical variables season (factor levels = fall, spring), burned level (unburned, moderately, heavily), submergence (traversing, lower part, upper part), decay (sound, decayed), form (straight, bent), position (ramp/bridge, loose), and stream order (first, second, third). The numeric fixed covariates were diameter, length, percentage within bankfull, number of anchor ends, distance to bank, and wood complexity. Because data records were nested by wood piece, which in turn were nested within reaches, we considered using both as random factors following that hierarchical order. However, the expected log-predictive density leave-one-out cross-validation difference (Δ ELPD LOO) suggested adding reach to random effects worsened the expected predictive accuracy (Δ ELPD LOO = −32.1 ± 8.5 SE). Prior to analysis, we centered and standardized the numeric covariates. A matrix of Spearman’s correlations for initial explanatory variables revealed no collinearity (|r_s_| ≤ 0.52 in all cases).

The minimal adequate (optimal) model was arrived at by first fitting the full model (with the 13 explanatory variables simultaneously) followed by backward elimination of one explanatory variable at a time. We used Watanabe-Akaike information criterion (WAIC; Watanabe 2010) to compare the relative fit of computed models to the data, interpreting WAIC differences greater than twice its corresponding standard error as suggesting that the model with the lower WAIC fitted the data substantially better. Using the WAIC criterion, distance to bank, wood complexity, and stream order were dropped in that order to reach the optimal model.

We created the Bayesian models in Stan computational framework (http://mc-stan.org/) accessed with *brms* package (Bürkner 2017). To improve convergence while controlling against overfitting, we assigned weakly informative priors to all the effect size beta parameters of the model (see Gelman 2020). We used the *normal (0, 10)* distribution for the beta in all levels of categorical variables except for season and the *normal (0, 5)* distribution for the beta in season and in the numeric variables. For each model, we ran four parallel MCMC chains until convergence was reached (all Rhat ≤ 1.1). Each chain had 4000 iterations (warmup = 1000, thin = 1), totaling 12,000 post-warmup samples. We assessed model adequacy using posterior predictive checks. We performed all analyses in R v. 3.6.3 (R Core Team 2020).

## Results

Over the three years of postfire wood function observations, we collected 1471 records for 567 pieces of wood. About 43% of the records were collected from burned wood (248 moderately and 385 heavily burned). The number of records was well balanced between fall (n = 700) and spring (771). Retention (385) and Substrate (368) were the most common functions, followed by Jam (223) and Flow (93). No function was observed in 402 records (Table 2).

**Table 2.**
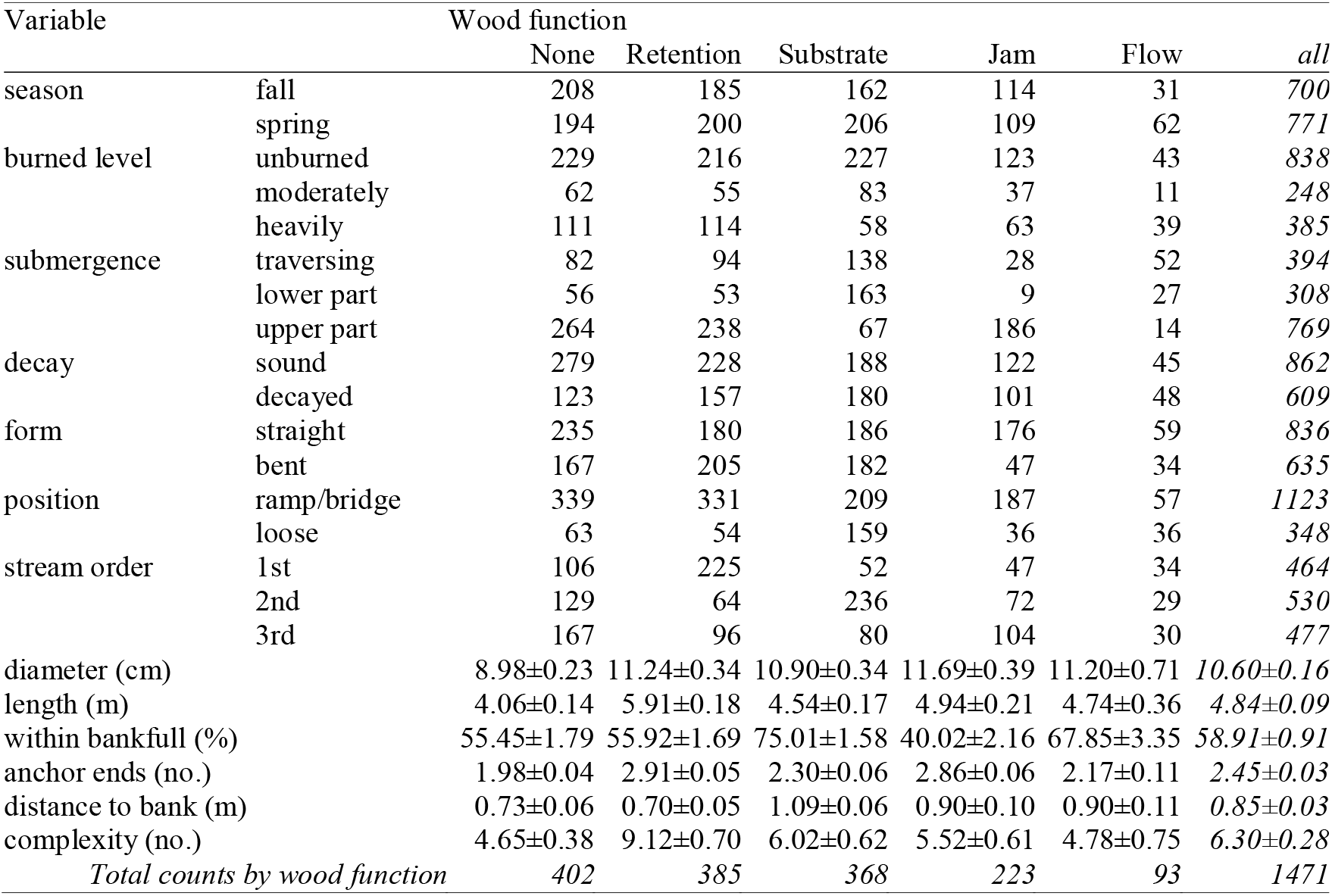
Variable summary by stream wood function, including counts by level in categorical variables and mean ± SE in numerical variables.

### Burned level and seasonal effects

When we accounted for the effect of the random factor (i.e., wood piece), our optimal mixed model (Table 3) showed that heavily burned wood was less likely to function as substrate for stream biota than unburned or moderately burned wood. After performing nonlinear hypothesis testing (Bürkner 2017, Clark 2020) for contrast effects between unburned, moderately, and heavily burned wood regarding the Substrate function, we are 100 % confident that heavily burned wood was less likely to provide substrate than each of the other two burn levels. The mean of the posterior distribution was 0.05 probability of heavily burned wood acting as a Substrate (95 % credible interval = 0.01–0.14), whereas this probability was 0.20 (0.09–0.37) and 0.31 (0.13–0.57) for unburned and moderately burned wood, respectively (Fig. 3). Thus, the probability of becoming Substrate was 4.0 and 6.2-fold lower in heavily burned wood than in unburned or moderately burned wood.

**Figure 3.**
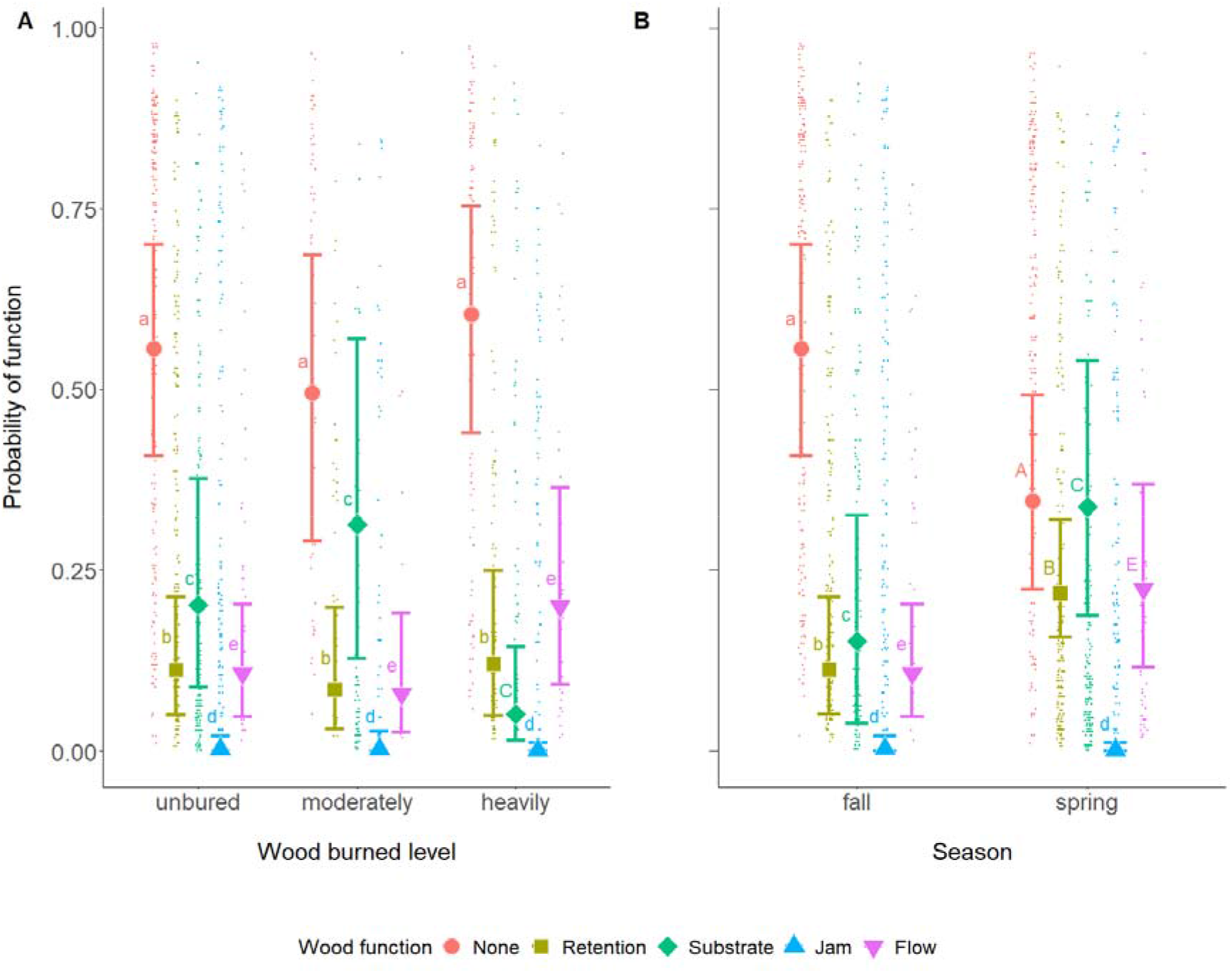
Mean fitted values (± 95% credible intervals) by stream wood burned level (A) and season (B) for the optimal multinomial logistic mixed-effects Bayesian model predicting the effects of these and other covariates on the main observable functions of stream wood. Small dots are predicted values. Different letter cases between levels of the same variable denote greater/lesser effects under the 95 % credible interval. None = stream wood without observable function; Retention = retaining organic matter; Substrate = serving as a substrate for aquatic biota; Jam = creating debris jams; Flow = deflecting flow (e.g., creating pools or riffles, forming steps).

**Table 3.**
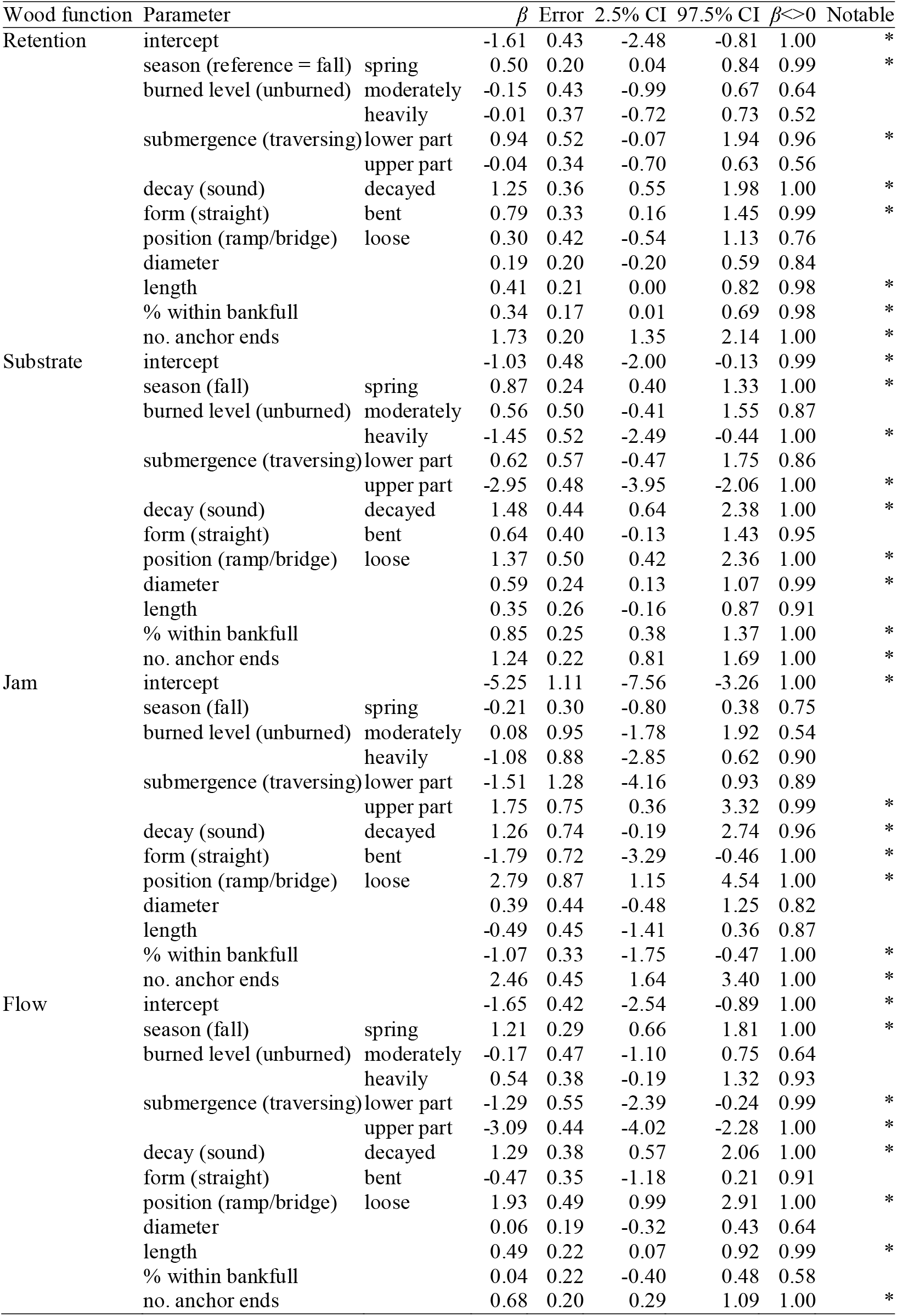
Summary of the fixed part of the multinomial mixed-effects Bayesian model predicting the effects of wood burned level, season, and covariates on the main observable functions of stream wood. Retention = retaining organic matter such; Substrate = serving as a substrate for aquatic biota; Jam = creating debris jams; Flow = deflecting flow (e.g., creating pools or riffles, forming steps). Non-observable function was the reference category. CI = credible interval for the parameter; β<>0 = posterior probability under the hypothesis of whether effect is greater (less) than zero if positive (negative); Notable = asterisk on effects whose CI does not contain zero (or with margin very close to zero). Potential scale reduction factor on split chains (Rhat) was 1.00 in all parameters.

As expected, the probability of each wood function in streams was contrasting between fall and spring. In spring, Retention (0.22; 95%-CI = 0.16–0.32), Substrate (0.34; 0.19–0.54), and Flow (0.22; 0.12–0.37) probabilities were 2.0, 2.3, and 2.0 times higher compared to the fall.

Posterior probabilities under the hypotheses testing that these effects were greater than zero (Clark 2020) allowed us to be 99 (Retention wood function) and 100 % (Substrate, Flow) confident that the probabilities of these functions were higher in the spring than in the fall. On the contrary, with regard to Jam function, our analysis allows us to be only 75 % confident that this function would appear less likely in the spring (0.001; 95%-CI: 0.00–0.01) compared to the fall (0.003; 0.00–0.02). The probability of non-observable function was 1.6 times higher in the fall (0.56; 95%-CI: 0.41–0.70) compared to spring (0.35; 95%-CI: 0.22–0.49).

### Notable covariate effects

Submergence had notable (|β-95%-CI| > ~0; Table 3) effects on the four functions, With the wood in the upper half of the channel decreasing its probability of serving as Substrate and deflecting Flow, while also decreasing the probability of forming Jams. Wood submerged in the lower half of the channel (lower half of bankfull height) favored the Retention (retaining organic matter such as twigs, leaves, fine organic matter) and Flow functions. Increased decay of wood positively influenced the four functions. As for form, relative to straight wood, bent wood increased the probability of Retention function and decreased the probability of Jam (creating debris jams) function. Wood position in the stream was an important factor, with loose wood (resting entirely on the streambed), relative to bridges and ramps, notably increasing the probability of all functions except Retention. As for wood size, diameter and length were contrasting for different functions; while diameter positively influenced the Substrate function, longer lengths favored the Retention and Flow functions. Relative to a piece traversing the bankfull height, the higher the percentage of the wood piece within the bankfull channel, the more likely it was to favor the Retention and Substrate functions. Pieces in the upper or lower halves were less likely to deflect the flow than those traversing the channel. Lastly, the probability of each of the four functions increased notably with the wood’s anchoring degree (number of anchor ends).

## Discussion

Through an extensive three-year seasonal tracking of stream wood following forest fires, we have produced empirical evidence that fire and season can alter the likelihood of ecologically important wood functions, especially when there is such a marked contrast between dry and wet seasons as in the Mediterranean. In comparable ecosystems, the success of postfire restoration efforts to offset the long-term impacts of forest wildfires in lotic ecosystems will benefit from our result that wood burned level can affect specific functions. In a fire-prone region with marked seasonality, we have also demonstrated that the probability of ecologically important functions follows seasonality. This intra-annual dynamic had not been examined previously. We expect our results will help guide postfire restoration efforts in comparable freshwater ecosystems as changing global climate and ongoing anthropogenic activities combine to increase the frequency and severity of fire around the world (Moreira et al. 2011, Coogan et al. 2019).

### Effect of wood burning level on its functions in streams

Our results strongly support that one of the main ecological functions of wood in world rivers, i.e. to provide substrate for biological organisms (Tank & Webster 1998, Dossi et al. 2020), can be negatively affected in heavily burned wood. This result expands upon another study investigating effects on the same functions, which did not establish a significant direct relationship with wood burn status, most likely because it considered the functions lumped together and reduced to simple binary criteria — with/without function (Vaz et al. 2013b). Our approach suggests that substrate provisioning by wood may be especially affected in the future, considering the growing global trend for more intense and severe wildfires (Moreira et al. 2011, Coogan et al. 2019) that are more likely to yield heavily burned wood (Agee 1993). Larger proportions of heavily burned wood serving less as substrate for vegetation, periphyton, biofilm, and ovipositions can negatively affect many xylobiont species and ultimately have implications for the response of the entire stream food web.

The information we collected on individual wood pieces was substantial, though our study could have been strengthened had we been able to include other basins for comparison. For example, although we did not detect an effect of stream order on wood functions, this could have changed had we considered more stream reaches with similar and larger sizes in more basins.

### Seasonal effects on stream wood functions

Our expectation that the probability of each stream wood function would change intra-annually was confirmed for three of the four functions studied and the probability of non-observable function was also nearly twice as high in the fall as in the spring. Although our study would benefit from including more sampling years, these results are novel and largely congruent with previous research showing the importance of season in the dynamics of ecological processes in non-perennial streams (García-Roger et al. 2011, Hodges & Magoulick 2011, Verkaik et al. 2013, Senter et al. 2017). None of these previous studies, however, documented the seasonal variation in the probability of each function for individual stream wood pieces. It is noteworthy that a function can be crucial even when it is less likely.

For example, our results showed, as expected, that flow deflection — including pool habitat formation — was less likely in the dry season relative to spring. Notwithstanding, it is also during that critical period that pool formation is paramount to increase the availability of food or refugia at the habitat scale in non-perennial Mediterranean rivers (Pires et al. 2010, Howell et al. 2012). As for jam formation, we found it surprising that it was less likely in the wet season unlike the other functions, though this result was more uncertain. One possibility was that detecting key pieces forming jams may have been slightly easier in dry seasons.

### Covariate effects of stream wood functions

Our data have comprehensively shown that varying primary metrics applied to stream wood (Wohl et al. 2010) can drive specific functions along with the burning level. Among wood relationships with the stream channel, our results showed that submergence and anchoring are factors clearly affecting the probability of each of the four functions following wildfires. Interestingly, we found different submergence levels to result in positive or negative contributions depending on the function, whereas greater anchoring was bound to increase the probability of all the functions examined. Thus, an eventual manipulation of stream wood submergence must take into account the function to be promoted. For instance, considering that heavily burned wood acts less as substrate, according to our results, that may be compensated by the presence of wood in the lower half of the channel after the fire. As for structural attributes, greater decay also has proven to be relevant in fostering the four wood functions as expected. This result is in line with previous research documenting decayed stream wood contributing more to matter retention, forming jams, bank stability, and riffle and pool formation (Gurnell et al. 1995, Jones et al., 2011). Possibly, pieces that have been downed wood for a longer time (i.e., with greater decay) have had more opportunity to become situated in stable locations where they are more likely to provide functions. Our data also highlight that some widely used metrics seemed to affect only certain functions. Although the “size paradigm” (Vaz et al. 2013b) assumes larger-sized pieces to provide more function, our results go further by suggesting the effect of wood size to be much more function-specific. For instance, the wood length was only more likely to have a notable positive effect on the probability of flow deflection.

### Management implications

Our study clearly demonstrates that wood burned level matters for the probability of its specific functions in the study streams within burnt areas and therefore managers must account for it in future postfire restoration campaigns. Given the high recruitment of wood following forest wildfires (Vaz et al. 2015) and the controversial impetus to remove some of it (e.g., to eliminate stream blockage), our study suggests the selective removal of the most heavily carbonized wood would allow wood to persist with the greatest potential to provide substrate for the stream biota. Nonetheless, even heavily burned wood can provide structural complexity and cover for fish, both of vital importance in sand-bed streams for instance (Gurnell et al. 1995). If wood must be removed postfire for safety, political, or administrative reasons, the burning level should not be the only criteria in choosing pieces to remove. For example, the relative non-attraction of heavily burned wood as substrate can be compensated for by other wood in the system with characteristics enhancing this function, such as wood deeper within the bankfull area, and with large diameters. If the objective is to choose nonfunctional wood, our raw data suggested pieces slightly smaller, more loosely anchored and less complex are better options. If the objective is to prevent postfire wood from moving downstream, a reasonable approach would be to remove pieces most likely to be mobilized, those that are not buried, submerged deep in the channel, relatively small, not braced, and lack rootwads (Merten et al. 2010).

Managers should also give consideration to adding unburned wood to streams postfire. Wood additions will be particularly valuable in areas that burned with high intensity, leading to a triple damaging situation where most wood recruited to the stream is heavily burned, postfire hydrology favors wood export, and large unburned wood may not be recruited to the stream for decades. Adding unburned wood provides better substrate and bolsters other wood functions. If the goal is to reduce the loss of substrate function or reduce the damage from related fire impacts (e.g., to dampen postfire hydrographs or store postfire sediment) then wood additions should occur immediately after the fire. Alternatively, if the goal is to reduce the loss of overall wood function in the long term until riparian regrowth is sufficient to again recruit unburned wood, managers might consider waiting until postfire hydrographs have begun to recover and added wood is less likely to be exported.

The time frame of ecological restoration operations is particularly relevant when considering disturbed streams, whether as a result of natural or anthropogenic influences, and their recovery after disturbance (Gurnell et al. 1995). As such, we highlight one novelty of our study by showing that primary wood functions are not static in the lotic ecosystem and documenting its intra-annual dynamics in non-perennial Mediterranean streams. Because such seasonal effects are likely widespread in other systems, we advocate restorers elsewhere to undertake similar pilot tracking of stream wood functions prior to large-scale stream restoration campaigns and thus identifying function-season relationships for the functions to be promoted. The timing (and method) used for wood placement should then be appropriate to the functions to be promoted but also to the deep knowledge of the local ecology. For example, some adverse impacts can be avoided by scheduling work to avoid fish spawning and other environmentally sensitive periods (Anton et al. 2011).

Irrespective of wildfire impacts, our results can help guide the restoration of non-perennial Mediterranean streams with marked seasonal differences. Depending on season, the consequences of added wood can be highly relevant for stream biota. Namely, if we consider flow deflection, including pool formation in the most stressful conditions of the dry season, we suggest adding wood that is longer, anchored, and traversing the bankfull. Moreover, to optimize substrate provisioning, large diameters is important.

Broadly, our results must be integrated into the knowledge resulting from tens of thousands of projects in which wood has been used to enhance in-river habitat throughout the world for over a century (Bernhardt et al. 2005, Thompson et al. 2018). We provide guidance as to when and what stream wood to remove — provided it is indeed necessary — as well as to the characteristics of the wood to be kept or added after fires when considering the attribute-probability of function relationships here established. We hope the results help to inform successful management efforts, as is increasingly asked from the science of restoration ecology (Suding et al. 2015).

## Authors’ contributions

PGV, CTR, PP conceived and designed the experiment; PGV collected the data, analyzed the data, and led the writing; ECM advised on field methods. All authors contributed critically to the drafts and gave final approval for publication.

## Data Accessibility Statement

The data used in this study will be deposited in the Figshare data repository after publication.

